# Prompt rewetting of drained peatlands reduces climate warming despite methane emissions

**DOI:** 10.1101/748830

**Authors:** Anke Günther, Alexandra Barthelmes, Vytas Huth, Hans Joosten, Gerald Jurasinski, Franziska Koebsch, John Couwenberg

## Abstract

Peatlands are strategic areas for climate change mitigation because of their matchless carbon stocks. Drained peatlands release this carbon to the atmosphere as carbon dioxide (CO_2_). Peatland rewetting effectively stops these CO_2_ emissions, but also re-establishes the emission of methane (CH_4_).

Essentially, management must choose between CO_2_ emissions from drained or CH_4_ emissions from rewetted peatland. This choice must consider radiative effects and atmospheric lifetimes of both gases, with CO_2_ being a weak but persistent and CH_4_ a strong but short-lived greenhouse gas. The resulting climatic effects are, thus, strongly time-dependent. We used a radiative forcing model to compare forcing dynamics of global scenarios for future peatland management using areal data from the Global Peatland Database. Our results show that CH_4_ radiative forcing does not undermine the climate change mitigation potential of peatland rewetting. Instead, postponing rewetting increases the long-term warming effect of continued CO_2_ emissions. Warnings against CH_4_ emissions from rewetted peatlands are therefore unjustified and counterproductive.

## Introduction

Each year, drained peatlands worldwide emit ∼2 Gt carbon dioxide (CO_2_) by microbial peat oxidation or peat fires, causing ∼5 % of all anthropogenic greenhouse gas (GHG) emissions on only 0.3 % of the global land surface_□_^1^. A recent study states that the effect of emissions from drained peatlands in the period 2020–2100 may comprise 12–41 % of the remaining GHG emission budget for keeping global warming below +1.5 to +2 °C_□_^2^. Peatland rewetting has been identified as a cost-effective measure to curb emissions_□_^3^, but re-establishes the emission of methane (CH_4_). In light of the strong and not yet completely understood impact of CH_4_ on global warming_□_^4,5^ it may seem imprudent to knowingly create or restore an additional source. Furthermore, there is considerable uncertainty on emissions from rewetted peatlands and some studies have reported elevated emissions of CH_4_ compared to pristine peatlands_□_^6–9^_□_.

The trade-off between CH_4_ emissions with and CO_2_ emissions without rewetting is, however, not straightforward: CH_4_ has a much larger radiative efficiency than CO_2_ ^(10)^. Yet, the huge differences in atmospheric lifetime lead to strongly time-dependent climatic effects. Radiative forcing of long-term GHGs (in case of peatlands: CO_2_ and N_2_O) is determined by *cumulative* emissions, because they factually accumulate in the atmosphere. In contrast, radiative forcing of near-term climate forcers (in case of peatlands: CH_4_) depends on the contemporary emission *rate* multiplied with the atmospheric lifetime^10,11^, because resulting atmospheric concentrations quickly reach a steady state of (sustained) emission and decay. Meanwhile, common metrics like global warming potential (GWP) and its ‘sustained flux’ variants^11,12^ fail to account for temporal forcing dynamics. These different atmospheric dynamics are relevant for the question how the various management scenarios will influence global climate and whether a scenario will amplify or attenuate peak global warming, i.e. the maximum deviation in global surface temperatures relative to pre-industrial times. An amplification of peak warming increases the risk of reaching major tipping points in the Earth’s climate system^13,14^.

Here, we explore how the different lifetimes of CO_2_/N_2_O vs. CH_4_ play out when assessing options for peatland rewetting as a climate warming mitigation practice. We compare the following global scenarios:

- ‘Drain_More’: The area of drained peatland continues to increase from 2020 to 2100 at the same rate as between 1990 and 2017
- ‘No_Change’: The area of drained peatland remains at the 2018 level
- ‘Rewet_All_Now’: All drained peatlands are rewetted in the period 2020-2040
- ‘Rewet_Half_Now’: Half of all drained peatlands are rewetted in the period 2020-2040
- ‘Rewet_All_Later’: All drained peatlands are rewetted in the period 2050-2070

These scenarios represent extreme management options and exemplify the differences caused by timing and extent of rewetting. For our modeling exercise, we focus on the direct human-induced climatic effects and conservatively assume pristine peatlands to be climate-neutral. Further, we assume that the maximum peatland area to be drained during the 21^st^ century equals the area that is already drained in 2018 (505,680 km^2^, Global Peatland Database^15^) plus an additional ∼5,000 km^2^ per year (average net increase of drained peatland area between 1990 and 2017^16^). For all scenarios, we apply IPCC default emissions factors^17^. To compare the radiative forcing effects of the different GHGs, we use a simplified atmospheric perturbation model that has been shown to provide reliable estimates of the climatic effects of peatlands^18^ (see Methods).

## Results and Discussion

Rewetting of drained peatlands instantly leads to climatic benefits compared to keeping the *status quo* (Figure 1). In case of rewetting all drained peatlands (scenarios ‘Rewet_All_Now’ and ‘Rewet_All_Later’) the radiative forcing stops increasing followed by a slow decrease. Since the response of global temperature is lagging behind changes in total radiative forcing by 15-20 years^19^, peatlands should be rewetted as soon as possible to have most beneficial (cooling) effects during peak warming, which AR5 climate models expect to occur after ∼2060 with increasing probability towards the end of the century^20^ (Figure 1).

**Figure 1.**
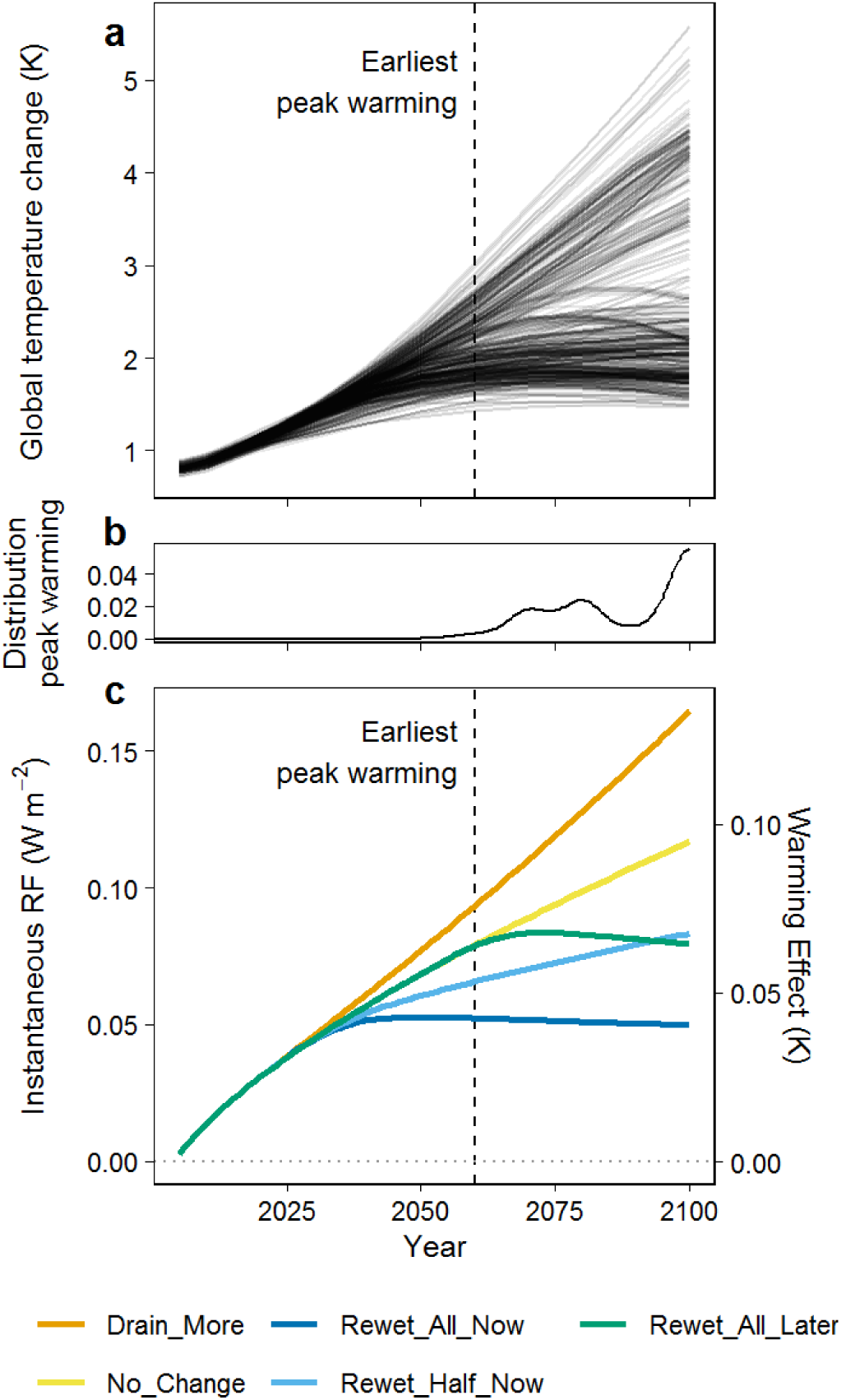
Climatic effects of peatland management in relation to global warming. Mean global temperature change relative to 2005 (a) and frequency distribution of the timing of peak warming (b) according to AR5 model pathways^20^ are shown compared to radiative forcings (RF) and estimated instantaneous warming effects of global peatland management scenarios (panel c, own calculations). Please note that in panel c) forcing of peatlands that remain pristine is assumed to be zero.

The overall climatic effect of peatland rewetting is indeed strongly determined by the radiative forcing of sustained CH_4_ emissions (Figure 2). However, because of the negligible or even negative emissions of CO_2_/N_2_O of rewetted peatlands and the short atmospheric lifetime of CH_4_, the total anthropogenic radiative forcing of all three GHGs combined quickly reaches a plateau after rewetting. Meanwhile, differences in radiative forcing between the ‘drainage’ (increased forcing) and ‘rewetting’ scenarios (stable forcing) are mainly determined by differences in the forcing of CO_2_ (Figure 2). Rewetting only half of the currently drained peatlands (‘Rewetting_Half_Now’) is not sufficient to stabilize radiative forcing. Instead, CO_2_ from not-rewetted peatland keeps accumulating in the atmosphere and warming the climate. Note that in the ‘Rewet_Half_Now’ scenario CH_4_ forcing is more than half that of the ‘Rewet_All_…’ scenarios, because drained peatlands also emit CH_4_, most notably from drainage ditches. Comparing the scenarios ‘Rewet_All_Now’ and ‘Rewet_All_Later’ shows that timing of peatland rewetting is not only important in relation to peak temperature, but also with respect to the total accumulated CO_2_ and N_2_O emissions in the atmosphere and the resulting radiative forcing (Figure 2).

**Figure 2.**
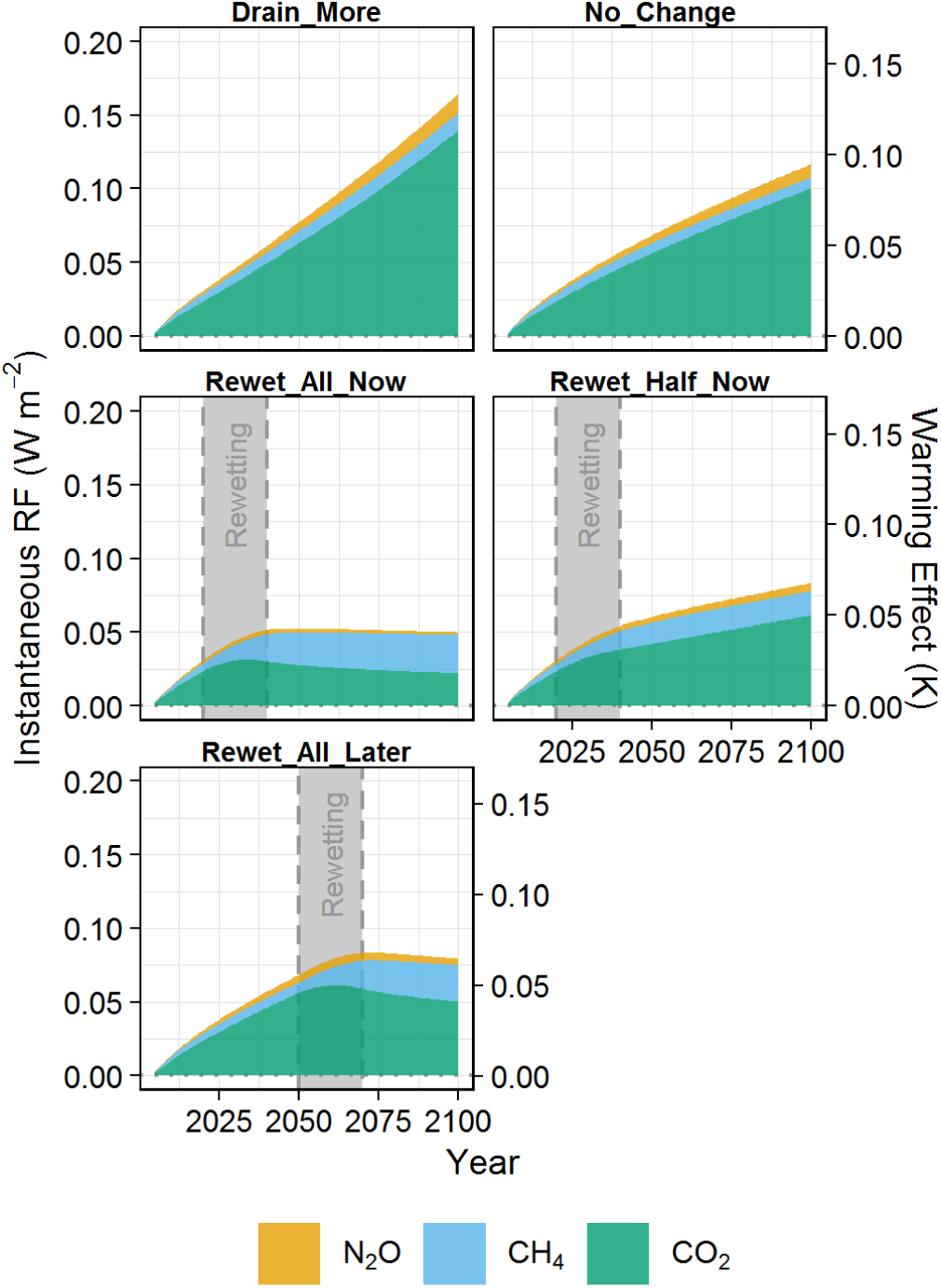
Contributions of the different GHGs (N_2_O, CH_4_, and CO_2_) to total radiative forcing (“RF”) and estimated warming effects in the modeled scenarios. The grey area shows the period of rewetting. Note that in the figure forcing of peatlands that remain pristine is assumed to be zero.

Our simulations highlight three general conclusions:

- The baseline or reference against which peatland rewetting has to be assessed is the drained state with its large CO_2_ emissions. For this reason, rewetted peatlands that are found to emit more CH_4_ than pristine ones_□_^9^ are no argument against rewetting. Moreover, whereas rewetted peatlands may again become CO_2_ sinks, the faster and larger climatic benefits of peatland rewetting result from the avoidance of CO_2_ emissions from drained peatlands.
- The climate effect is strongly dependent on the concrete point in time that rewetting is implemented. This fact is hitherto insufficiently recognized because it remains hidden by the common use of metrics that involve predetermined time horizons (like GWP or sustained flux variants of GWP).
- In order to reach climate-neutrality in 2050 as implied by the Paris Agreement, it is insufficient to focus rewetting efforts on selected peatlands only: to reach the Paris goal, CO_2_ emissions from (almost) all drained peatlands have to be stopped by rewetting_□_^2^.

Limiting global warming requires immediate reduction of global GHG emissions. It has been suggested that the negative climate effects of drained peatlands could be offset by growing highly-productive bioenergy crops^21^ or wood biomass^22^ as substitute for fossil fuels. In this study, we did not include this option because similar biomass-based substitution benefits can also be reached by cultivating biomass on rewetted peatlands^23^, i.e. without CO_2_ emissions from drained peat soil.

In conclusion, without rewetting the world’s drained peatlands will continue to emit CO_2_, with direct negative effects on the magnitude and timing of global warming. These effects include a higher risk of reaching tipping points in the global climate system and possible cascading effects^13^. In contrast, we show that peatland rewetting can be one important measure to reduce climate change and attenuate peak global warming: The sooner drained peatlands are rewetted, the better it is for the climate. Although the CH_4_ cost of rewetting may temporarily be substantial, the CO_2_ cost of inaction will be much higher.

## Methods

### Scenarios

Drained peatland area was taken from the Global Peatland Database (GPD)^15^, which includes *inter alia* national data from the most recent UNFCCC National Inventory Submissions and Nationally Determined Contributions. We used data separated by IPCC climate zone (boreal, temperate, and tropical) and assigned land use categories. Available land use categories were “Forest”, “Cropland”, “Deep-drained grassland”, “Shallow-drained grassland”, “Agriculture” (i.e. either grassland or cropland when the original data source did not differentiate between these two categories), and “Peat extraction” (see Table M1). Because of their only small area and uncertain emission factors, arctic drained peatlands (∼100 kha) were neglected. Newly drained/rewetted area in the scenarios is distributed across the climatic zones (and land use classes) according to the relative proportions of today’s drained peatland area. As future drainage – similar to the past two decades^16^ – will probably focus on tropical and subtropical peatlands, our ‘Drain_More’ scenario likely underestimates the climate effects of future drainage. For information on how variations in the assumed drainage rate and uncertainty of emission factors affected the displayed radiative forcing effects of the scenarios please see Fig. M1.

**Table M1.** Areas of drained peatland (kha) by climate zone and land use category according to the Global Peatland Database, together with aggregated emission factors. Emission factors assumed for rewetted peatlands are also shown for each climatic zone.

| Climatic zone | Land use category | Area<br>(kha) | CO<sub>2</sub><br>(t ha<sup>-1</sup> a<sup>-1</sup>) | CH<sub>4</sub><br>(kg ha<sup>-1</sup> a<sup>-1</sup>) | N<sub>2</sub>O<br>(kg ha<sup>-1</sup> a<sup>-1</sup>) |
| --- | --- | --- | --- | --- | --- |
| Boreal | Forest | 5474 | 2.5 | 9.8 | 2.6 |
|  | Cropland | 262 | 27.9 | 58.3 | 19.4 |
|  | Deep-drained grassland | 426 | 20.2 | 59.6 | 14.2 |
|  | Shallow-drained grassland | 0 | - | - | - |
|  | Agriculture | 3420 | 24.1 | 43.0 | 16.8 |
|  | Peat extraction | 333 | 10.2 | 32.9 | 0.5 |
|  | Rewetted | - | -1.3 | 123.6 | 0 |
| Temperate | Forest | 6315 | 10.3 | 7.9 | 4.3 |
|  | Cropland | 2528 | 28.6 | 58.3 | 19.4 |
|  | Deep-drained grassland | 3405 | 22.3 | 73.5 | 12.3 |
|  | Shallow-drained grassland | 2422 | 13.6 | 63.4 | 2.4 |
|  | Agriculture | 8389 | 21.0 | 55.8 | 10.1 |
|  | Peat extraction | 662 | 10.8 | 32.9 | 0.5 |
|  | Rewetted | - | -0.4 | 205.9 | 0 |
| Tropical | Forest | 7235 | 22.0 | 50.0 | 3.7 |
|  | Cropland | 305 | 45.0 | 118.9 | 4.2 |
|  | Deep-drained grassland | 70 | 37.4 | 52.0 | 7.7 |
|  | Shallow-drained grassland | 0 | - | - | - |
|  | Agriculture | 9314 | 42.5 | 96.6 | 5.4 |
|  | Peat extraction | 8 | 10.1 | 32.9 | 5.6 |
|  | Rewetted | - | 1.9 | 166.5 | 0 |

**Fig. M1.**
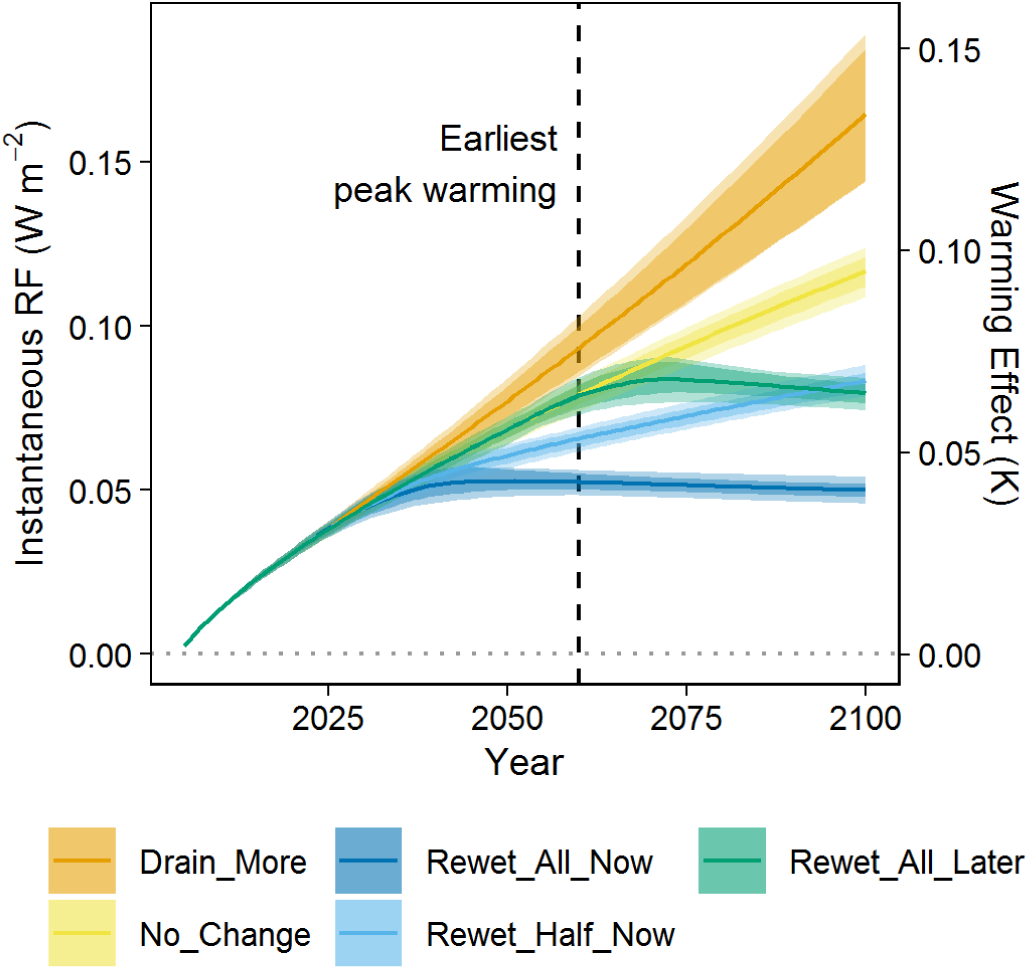
Sensitivity of radiative forcings (“RF”) and estimated warming effects of global peatland scenarios to modeling choices and uncertainty of emission factors. Error ranges represent the range of radiative forcing resulting from random variations in ongoing drainage rate (1000-8000 km^2^ per year) and IPCC emission factors (10 % and 20 % uncertainty of emission factor).

### Emissions

Emission factors for each climate zone and land use category were taken from the IPCC Wetland supplement^17^ that presents the most robust and complete meta-study of published emission data. Emission factors were averaged for IPCC categories that were given at a higher level of detail (e.g. nutrient-poor vs. nutrient-rich boreal forest) than the available land use categories from the GPD. Equally, we averaged the supplied emission factors for grassland and cropland in order to obtain emission factors of the land use class “Agriculture” (see Table M1 for final aggregated emission factors and Supplementary Table S1 for exact aggregation steps). We included emissions from ditches and DOC exports by using emission factors and default cover fraction of ditches given by the IPCC^17^ (Table S1). Since the IPCC Wetlands Supplement does not provide an emission factor for CH_4_ from tropical peat extraction sites, we assumed the same CH_4_ emissions as for temperate/boreal peat extraction. Values of the emission factors could change slightly when more emission data becomes available. To cover this possibility, we randomly varied all emission factors within a range of 10 % and 20 % uncertainty in our sensitivity analysis (Fig. M1). Individual studies have discussed the presence of a CH_4_ peak for the first years after rewetting_□_^7,8^. Although this is likely not a global phenomenon^24^, please see supplementary Figure S1 for an estimate of the uncertainty related to possible CH_4_ peaks.

### Radiative forcing

The forcing model uses simple impulse-response functions^25^ to estimate radiative forcing effects of atmospheric perturbations of CO_2_, CH_4_ and N_2_O fluxes^12^. Perturbations of CH_4_ and N_2_O were modeled as simple exponential decays, while CO_2_ equilibrates with a total of five different pools at differing speeds. For CO_2_, we adopted the flux fractions and perturbation lifetimes used by ref^18^. In the model, we assume a perfectly mixed atmosphere without any feedback mechanisms but include indirect effects of CH_4_ on other reagents and aerosols^10^.

Climatic effects of CO_2_ from CH_4_ oxidation should not be considered for CH_4_ from biogenic sources^10^. However, although the large majority of CH_4_ from peatlands stems from recent plant material (a biogenic source), the proportion of fossil CH_4_ (from old peat) may be substantial in some cases^26^. Thus, we conservatively included the climatic effect of CO_2_ from CH_4_ oxidation in our analyses. Overall, this forcing comprised only 5-7 % of the CH_4_ radiative forcing and only ∼1-3 % of total radiative forcing.

We compare the radiative forcing trajectories of the various peatland management scenarios with the global temperature change as projected by all available pathways of IPCC’s AR5^20^ and use the same starting year 2005 as these pathways. Further, we estimated the approximate effects of radiative forcing on global mean temperature as ∼1 K per 1.23 W/m^2^ radiative forcing^27^.

### Data availability

The models for projected temperature change were downloaded from the stated website. Emission factors and peatland cover data are entirely included in the manuscript. The code for the atmospheric perturbation model can be found in the supplementary information.

## Acknowledgements

The European Social Fund (ESF) and the Ministry of Education, Science and Culture of Mecklenburg-Western Pomerania funded this work within the scope of the project WETSCAPES (ESF/14-BM-A55-0030/16 and ESF/14-BM-A55-0031/16). G. J. received funding within the framework of the Research Training Group Baltic TRANSCOAST from the DFG (Deutsche Forschungsgemeinschaft) under grant number GRK 2000/1. This is Baltic TRANSCOAST publication no. GRK2000/00XX. V. H. gratefully acknowledges funding by the Federal Agency of Nature Conservation (BfN, grant number: 3516892003) and by the European Regional Development Fund (ERDF) distributed through the NBank.

## Author contributions

A. G., J. C., G. J. and V.H. conceived the study. A. G., A. B., J. C., H. J. assembled input data. A. G. implemented the simulation model with contributions from J. C. All authors discussed the results and implications. A. G. led writing of the manuscript with comments/edits from all authors.

## Author information

The authors declare no competing interests.

