## Supplemental Figure S1 and Table S1 for "Prompt rewetting of drained peatlands reduces climate warming despite methane emissions"

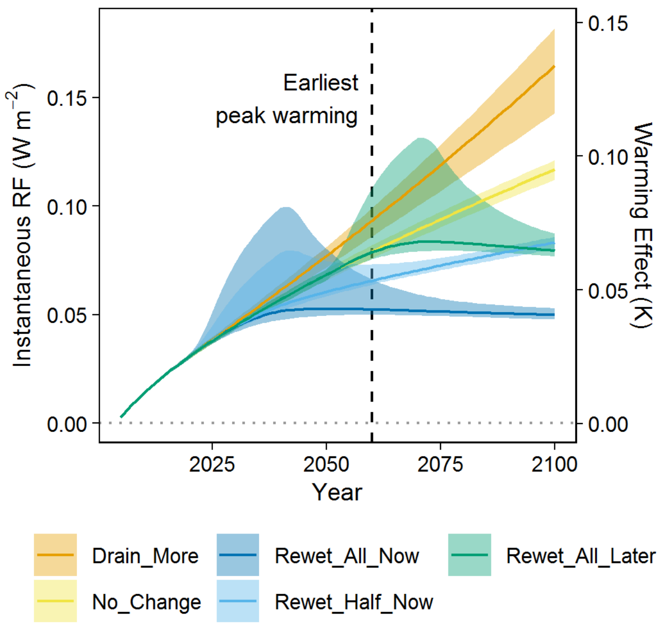


Figure S1 Sensitivity of radiative forcings ("RF") and estimated warming effects of global peatland scenarios to modeling choices, uncertainty of emission factors and possibility of methane peaks after rewetting. Error ranges represent the range of radiative forcing resulting from random variations in ongoing drainage rate (1000-8000 km² per year), IPCC emission factors (10 % and 20 % uncertainty of emission factor), and occurrence of a methane peak (10 times rewetted methane emission factor for 5 years after rewetting).

Table S1 Assigned and aggregated IPCC emission factors (Wetlands Supplement) of GPD categories.

| GPD | | IPCC | | | | | | |
|---|---|---|---|---|---|---|---|---|
| | | Field | | $CO_2$ from DOC | | Ditches Cover fraction | $CH_4$ | |
| Climatic zone | Land use category | Climatic zone | EF | Climatic zone | EF | | Climatic zone | EF |
| Boreal | Forest | Boreal | Average of 'Drained Forest Land Nutrient-poor' and 'Drained Forest Land Nutrient-rich' | Boreal | 'Drained' | 2.5% | Boreal / Temperate | 'Drained Forest Land / Drained Wetlands' |
| Boreal | Cropland | Boreal / Temperate | 'Drained Cropland' | Boreal | 'Drained' | 5.0% | Boreal / Temperate | 'Deep-drained Grassland / Cropland' |
| Boreal | Deep drained grassland | Boreal | 'Drained Grassland' | Boreal | 'Drained' | 5.0% | Boreal / Temperate | 'Deep-drained Grassland / Cropland' |
| Boreal | Agriculture | Boreal | Average of 'Drained Cropland' and 'Drained Grassland' | Boreal | 'Drained' | 5.0% | Boreal / Temperate | Average of 'Deep-drained Grassland / Cropland' and 'Shallow-drained Grassland' |
| Boreal | Peat extraction | Boreal / Temperate | 'Peatland Managed for Extraction' | Boreal | 'Drained' | 5.0% | Boreal / Temperate | 'Peat Extraction' |
| Boreal | Rewetted | Boreal | Average of 'Rewetted Poor' and 'Rewetted Rich' | Boreal | 'Rewetted' | 5.0% | Boreal / Temperate | 'Drained Wetland / Forest' |
| Temperate | Forest | Temperate | 'Drained Forest Land' | Temperate | 'Drained' | 2.5% | Boreal / Temperate | 'Drained Forest Land / Drained Wetlands' |
| Temperate | Cropland | Boreal / Temperate | 'Drained Cropland' | Temperate | 'Drained' | 5.0% | Boreal / Temperate | 'Deep-drained Grassland / Cropland' |
| Temperate | Deep drained grassland | Temperate | Average of 'Deep-drained Nutrient-rich Grassland' and 'Drained Nutrient-poor Grassland' | Temperate | 'Drained' | 5.0% | Boreal / Temperate | 'Deep-drained Grassland / Cropland' |
| Temperate | Shallow-drained grassland | Temperate | Average of 'Shallow-drained Nutrient-rich Grassland' and 'Drained Nutrient-poor Grassland' | Temperate | 'Drained' | 5.0% | Boreal / Temperate | 'Shallow-drained Grassland' |
| Temperate | Agriculture | Temperate | Average of 'Drained Cropland', 'Deep-drained Nutrient-rich Grassland', 'Shallow-drained Nutrient-rich Grassland' and 'Drained Nutrient- | Temperate | 'Drained' | 5.0% | Boreal / Temperate | Average of 'Deep-drained Grassland / Cropland' and 'Shallow-drained Grassland' |

| | | | | | | | | |
|---|---|---|---|---|---|---|---|---|
| | | | poor Grassland' | | | | | |
| Temperate | Peat extraction | Boreal / Temperate | 'Peatland Managed for Extraction' | Temperate | 'Drained' | 5.0% | Boreal / Temperate | 'Peat Extraction' |
| Temperate | Rewetted | Temperate | Average of 'Rewetted Poor' and 'Rewetted Rich' | Temperate | 'Rewetted' | 5.0% | Boreal / Temperate | 'Drained Wetland / Forest' |
| Tropical | Forest | Tropical | 'Drained Forest Land' | Tropical | 'Drained' | 2.0% | Tropical | 'Drained' |
| Tropical | Cropland | Tropical | Average of 'Drained Cropland and Fallow' and 'Drained Cropland: Paddy Rice' | Tropical | 'Drained' | 2.0% | Tropical | 'Drained' |
| Tropical | Deep drained grassland | Tropical | 'Drained Grassland' | Tropical | 'Drained' | 2.0% | Tropical | 'Drained' |
| Tropical | Agriculture | Tropical | Average of 'Drained Cropland and Fallow', 'Drained Cropland: Paddy Rice' and 'Drained Grassland' | Tropical | 'Drained' | 2.0% | Tropical | 'Drained' |
| Tropical | Peat extraction | $CO_2$/$N_2O$: Tropical, $CH_4$: Boreal / Temperate | 'Peatland Managed for Extraction' | Tropical | 'Drained' | 2.0% | Tropical | 'Drained' |
| Tropical | Rewetted | Tropical | 'Rewetted' | Tropical | 'Rewetted' | 2.0% | Tropical | 'Drained' |
